# Mapping and dynamics of regulatory DNA in maturing seeds

**DOI:** 10.1101/235325

**Authors:** Alessandra M. Sullivan, Andrej A. Arsovski, Agnieszka Thompson, Richard Sandstrom, Robert E. Thurman, Shane Neph, Audra K. Johnson, Shawn T. Sullivan, Peter J. Sabo, Fidencio V. Neri, Molly Weaver, Morgan Diegel, Jennifer L. Nemhauser, John A. Stamatoyannopoulos, Kerry L. Bubb, Christine Queitsch

## Abstract

The genome is reprogrammed during development to produce diverse cell types, largely through altered expression and activity of key transcription factors. The accessibility and critical functions of epidermal cells have made them a model for connecting transcriptional events to development in a range of model systems. In *Arabidopsis thaliana* and many other plants, fertilization triggers differentiation of specialized epidermal seed coat cells that have a unique morphology caused by large extracellular deposits of pectin. Here, we used DNase I-seq to generate regulatory landscapes of *A. thaliana* seeds at two critical time points in seed coat maturation, enriching for seed coat cells with the INTACT method. We found over 3000 developmentally dynamic regulatory DNA elements and explored their relationship with nearby gene expression. The dynamic regulatory elements were enriched for motifs for several transcription factors families; most notably the TCP family at the earlier time point and the MYB family at the later one. To assess the extent to which the observed regulatory sites in seeds added to previously known regulatory sites in *A. thaliana*, we compared our data to 11 other data sets generated with seven-day-old seedlings for diverse tissues and conditions. Surprisingly, over a quarter of the regulatory, *i.e.* accessible, bases observed in seeds were novel. Notably, in this comparison, development exerted a stronger effect on the plant regulatory landscape than extreme environmental perturbations, highlighting the importance of extending studies of regulatory landscapes to other tissues and cell types during development.

## Introduction

Spatial and temporal regulation of gene expression is critical for development and specialization of tissues and cell types. *Cis*-regulatory DNA elements, and the *trans*-acting factors that bind them, are a primary mechanism for regulating gene expression. Active *cis*-regulatory elements such as promoters, enhancers, insulators, silencers, and locus control regions can be identified by their characteristic hypersensitivity to cleavage by DNase I (Banerji, Olson, and Schaffner 1983; Baniahmad et al. 1990; Chung, Bell, and Felsenfeld 1997; Talbot et al. 1989; Thurman et al. 2012; Wu et al. 1979; Wu, Wong, and Elgin 1979). Our previous analyses of regulatory DNA and its dynamics in *A. thaliana* largely focused on identifying regulatory networks and divergence of regulatory DNA in whole seedlings (A. M. Sullivan et al. 2014). Our method, which relies on INTACT-labeled nuclei (Deal and Henikoff 2010), lends itself to investigating the regulatory landscape of nuclei enriched for certain cell types. Cell-type-enriched, and ideally cell-type-specific, approaches to gene regulation and expression are fundamental for understanding development. Here, we use DNase I-seq to examine the regulatory landscape of seeds at two critical developmental time points, four and seven days post-anthesis, enriching for seed coat cells as they transition from the non-mucous-secreting state to the mucous-secreting state.

The seed coat differentiates from the integuments of the ovule after fertilization has occurred. In many species, seed coat cells produce and store polysaccharide-rich mucilage (myxospermy). When wetted, this mucilage expands and extrudes from mucous-secreting cells, forming a gel-like layer around the seed (Western, Skinner, and Haughn 2000; Windsor et al. 2000). In *A. thaliana*, mucilage is composed primarily of pectin with lesser amounts of cellulose and xyloglucan (Haughn and Western 2012). Although the function of mucilage depends on the species and the environmental context (García-Fayos, Bochet, and Cerdà 2010; Garwood 1985; Gutterman and Shem-Tov 1997; Yang, Dong, and Huang 2010; Yang et al. 2011), mucilage is generally thought to protect the emerging seedling and facilitate its germination.

In *A. thaliana*, seed coat cell differentiation and maturation is well characterized at the morphological level (Western, Skinner, and Haughn 2000; Windsor et al. 2000). In the mature ovule, seed coat cells contain a large vacuole. During the first four days after fertilization, the vacuole swells causing cell growth, and starch granules appear. By seven days after fertilization, the vacuole shrinks, the cytoplasm forms a column filled with vesicles and golgi stacks, and mucilage is secreted into the apoplast. By ten days post fertilization, mucilage production is complete, and a secondary cell wall is being deposited around the columnar cytoplasm forming a solid structure, the columella. Once differentiation is complete, dehydration shrinks the stored mucilage causing the primary cell wall to drape over the newly formed columella, creating the polygonal pattern visible on the dry seed exterior.

Seed coat cells are an exceptionally well-studied plant cell type. Previous studies have identified 48 genes affecting seed coat cell differentiation and maturation when disrupted in *A. thaliana* (Francoz et al. 2015; North et al. 2014). These genes fall into roughly three functional categories: epidermal cell differentiation, mucilage synthesis and secretion, and secondary cell wall synthesis (**Supplemental Table 1**). Genes controlling specification of the ovule integument will also impact seed coat cell differentiation. Many of the genes required for seed coat differentiation and mucilage production are transcription factors (**Supplemental Table 1**) (Francoz et al. 2015).

While the identity of the TFs, and in some cases their targets, are known, there is little information about individual regulatory elements and their activity during seed coat differentiation and maturation. Exceptions include the promoter of *DP1*, which specifically drives seed coat epidermal expression (Esfandiari et al. 2013), and the L1 box in the *CESA5* promoter, which interacts with GL2 (a seed coat epidermis differentiation factor) in yeast (Tominaga-Wada et al. 2009).

To address this paucity of genome-wide regulatory information, we employed the INTACT method to capture the nuclei of *GL2*-expressing cells from whole siliques, followed by DNase I-seq to identify regulatory elements, their dynamics, and their constituent TF motifs at two critical time points in seed development. We observe dramatic changes in the regulatory landscape, relate dynamic DNase I-hypersensitive sites (DHSs) to previously established expression profiles, identify genes that neighbor dynamic DHSs, and identify associated transcription factor motifs. We identify many candidate genes that may contribute to seed coat development in ways that might escape traditional genetic analysis.

By comparing our novel seed coat-enriched regulatory landscapes to previously generated landscapes we identified surprisingly many novel regulatory sites. Through this comparative analysis we also show that, like animals (Stergachis et al. 2013; Thomas et al. 2011; Daugherty et al. 2017), cell lineage and developmental stage are strong determinants of the plant chromatin landscape compared to even severe environmental perturbations. This result was somewhat unexpected given that plants are so exquisitely responsive to environmental cues. Taken together, our findings call for a systematic analysis of important *A. thaliana* cell types during development and in response to major environmental cues.

## Results

### The regulatory DNA landscape of maturing seed coat epidermal cells

To capture the regulatory landscape of seed coat epidermal cells, we employed nuclear capture (INTACT) (Deal and Henikoff 2010) followed by DNase I-seq (A. M. Sullivan et al. 2014). We used an existing transgenic plant line (Deal and Henikoff 2010) in which the *GL2* promoter controls the targeting of biotin to the nuclear envelope (**Supplemental Figure 1**). GL2 is expressed at very high levels in the seed coat epidermis; it is also expressed to varying degrees elsewhere in the seed, most noticeably in the embryo (Windsor et al. 2000; Belmonte et al. 2013). We sampled whole siliques, which encase 40-60 seeds, at 4 and 7 days post-anthesis (DPA), to capture the regulatory landscape before and after mucilage production begins in the seed coat.

We created five DNase I-seq libraries, including biological replicates for each time point, and identified a union set of 43,120 DHSs. Of these DHSs, 3,109 were determined to be developmentally dynamic between the 4DPA and 7DPA samples by DEseq2 (Love, Huber, and Anders 2014) with an adjusted p-value of < 0.001 (**Figure 1A, Supplemental Tables 2-5, Methods**). We denote DHS more accessible in 7DPA than 4DPA as activated DHSs, and those more accessible in 4DPA than 7DPA as deactivated DHSs.

**Figure 1.**
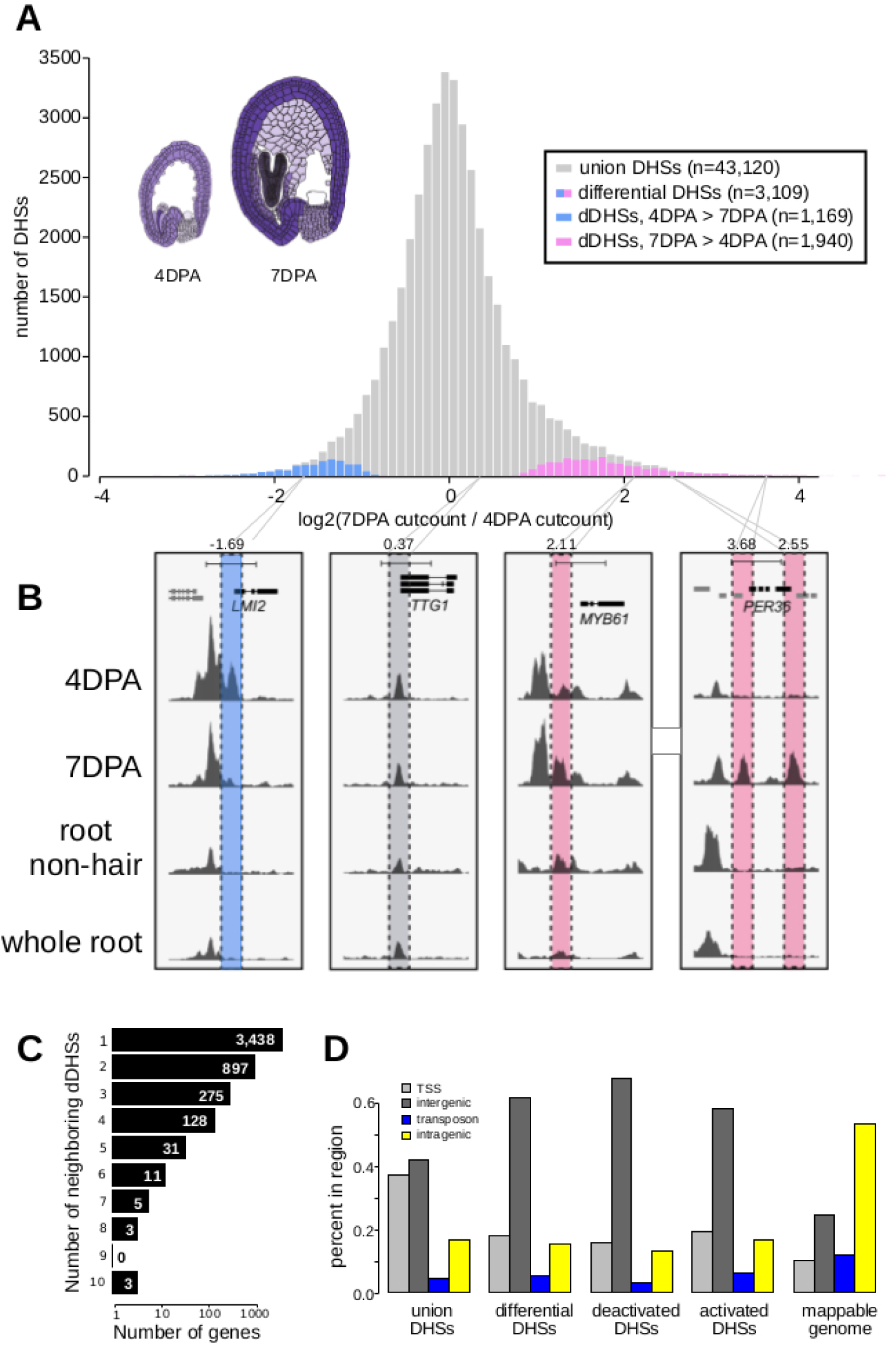
The chromatin landscape of maturing seed coat cells. **A**, Distribution of log2(DNase I cut count in 7DPA / DNase I cut count in 4DPA) for all union DHSs (gray) and differential DHSs, with DHSs more accessible at 4DPA appearing on the left in blue and DHSs more accessible at 7DPA appearing on the right in pink. Diagrams of 4DPA (left) and 7DPA seeds (right) are shown, with purple opacity indicating *GL2* expression levels from Belmonte et al. 2013. **B**, Examples showing a deactivated DHS, two examples of activated DHSs, and one example of a static DHS. A 5 kb region is shown in each window; all data tracks are read-depth normalized. **C**, Distribution of the number of dynamic DHSs neighboring genes. Most genes reside next to one dynamic DHS; however, surprisingly many genes reside next to multiple dynamic DHSs. **D**, The numbers of union DHSs (uDHSs) and dynamic DHSs (dDHSs) within each genomic context: TSS, intergenic, transposon, and intragenic.

Twenty activated DHSs resided near one of the 48 known seed coat development genes (**Supplemental Table 1)**, which represents a 2.5-fold enrichment over the eight genes expected by chance. For example, we found 7DPA-activated DHSs near *MYB61*, which is required for mucilage production (Penfield et al. 2001), and *PER36*, which is required for proper mucilage release (Kunieda et al. 2008) (**Figure 1B**). We also identified many dynamic DHSs near genes that were not previously associated with seed coat development. For example, the meristem identity transition transcription factor gene, *LMI2* (Pastore et al. 2011), resides near a DHS that was deactivated during seed coat cell maturation (**Figure 1B**). Similar to previous observations (A. M. Sullivan et al. 2014), the majority of observed DHSs were static during development, such as those flanking *CESA5*, which encodes a cellulose synthase that produces seed mucilage cellulose (S. Sullivan et al. 2011) (**Figure 1B**). The regulatory landscape of seed coat cells differed significantly from the landscape of root non-hair cells, another epidermal cell type, as well as from whole roots (**Figure 1B, Figure 5**). Consistent with multiple regulatory inputs in development, we observed that developmentally dynamic DHSs were frequently clustered with about a third of genes residing near more than one dynamic DHS (**Figure 1C**). We conclude our method detects developmentally regulated DHSs, which appear in the vicinity of known seed coat development genes and genes newly implicated in seed maturation.

Next, we asked whether the genomic distribution of dynamic DHSs was different than that of all DHSs by tabulating the number of DHSs occurring in various genomic contexts (*e.g.* intragenic) (**Supplemental Table 6**). Similar to whole seedling DHSs (A. M. Sullivan et al. 2014), DHSs in seed-coat-enriched cells (both dynamic and static), tended to reside in intergenic regions and near transcription start sites (TSSs, 400 bp upstream of the TSS), and were depleted in intragenic regions and transposable elements (TEs). In contrast, developmentally dynamic DHSs were primarily enriched in intergenic regions (**Figure 1D**). This distribution is consistent with previous observations in Drosophila, where developmental enhancers are primarily located in intergenic regions and in introns while housekeeping gene enhancers are primarily located near transcription start sites (Zabidi et al. 2015).

### Genes neighboring dynamic DHSs are enriched for differentially expressed genes

Of the 28,775 annotated genes in TAIR10, 4,791 (16.6%) neighbor one or more of the 3,109 developmentally dynamic DHSs, with a few genes flanked by as many as ten developmentally dynamic DHS (**Figure 1C**). As we and others have shown previously, chromatin accessibility is only weakly correlated with nearby gene expression (A. M. Sullivan et al. 2015); however, dynamic chromatin accessibility (*i.e.* dynamic DHSs) is more frequently correlated with altered expression of nearby genes. To explore the relationship between chromatin accessibility and gene expression in maturing seeds, we took advantage of two published seed coat epidermis expression studies (Belmonte et al. 2013; Dean et al. 2011), considering a gene to be differentially expressed if it exhibited a 2-fold expression change between developmental time points.

In the first study, Dean et al. 2011 quantified gene expression in manually dissected seed coats at 3DPA and 7DPA in the Col-2 accession, identifying 3,423 genes that exhibited at least a 2-fold expression change between these developmental stages (**Figure 2A, B, Supplemental Figure 2A, B)**. In the second study, Belmonte et al. 2013 quantified gene expression in many parts of the seed at many time points in the Ws-0 accession using laser capture micro dissection. For our analysis, we used the seed coat and embryo proper expression values from globular (∼3-4 DPA), heart (∼4-5 DPA) and linear cotyledon (∼7DPA) stage seeds; the former approximating the 4DPA stage while the latter approximates the 7DPA stage (Le et al. 2010). A total of 4,115 genes exhibited at least a 2-fold expression change in seed coat. Both studies used microarrays to evaluate gene expression (**Figure 2A, B, Supplemental Figure 2A, B)**.

**Figure 2.**
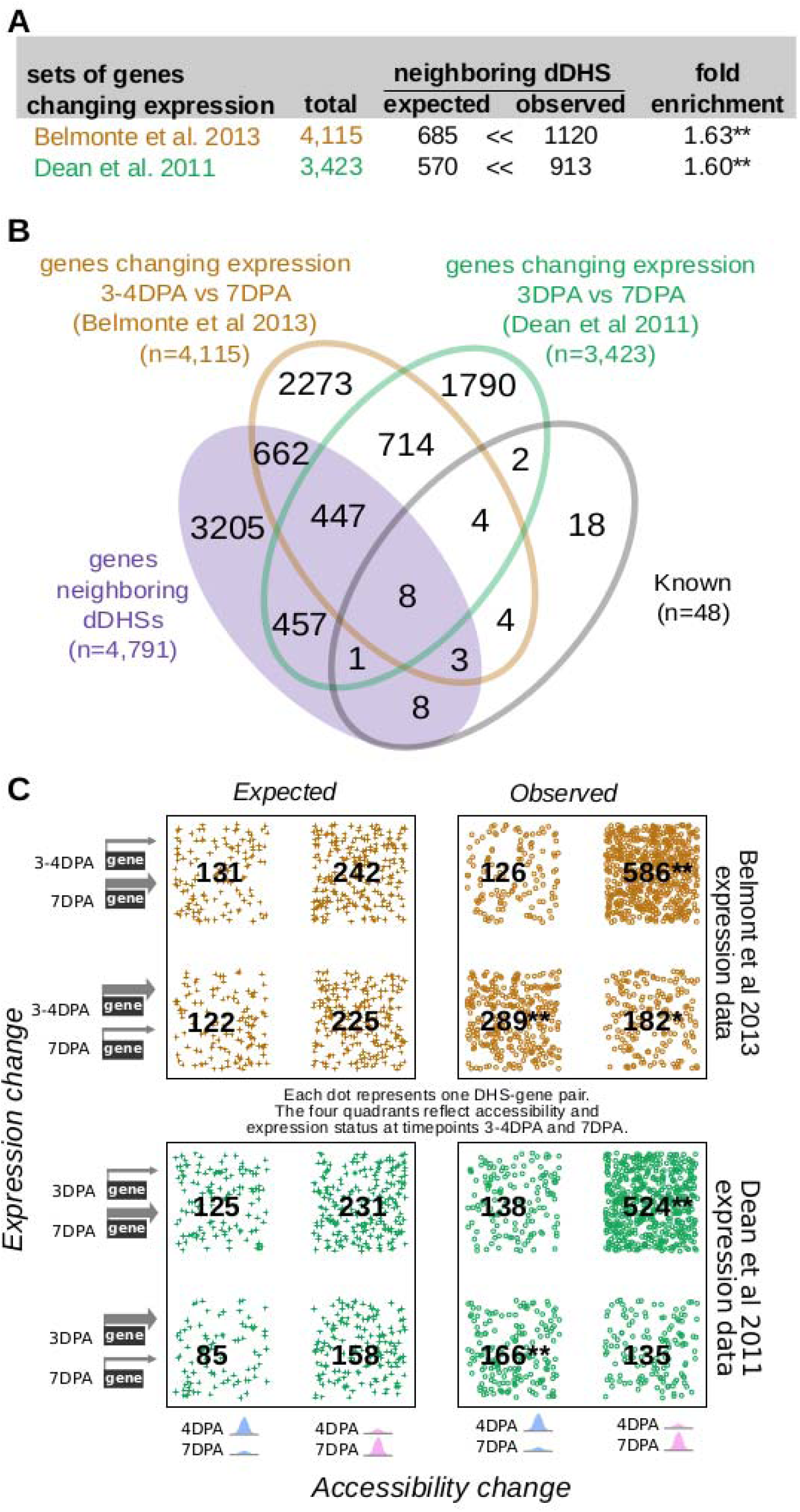
Genes neighboring developmentally dynamic DHSs are often differentially expressed. **A,** Overlap between the set of genes neighboring dDHSs and genes found to be differentially expressed in seed coat at stages 4DPA and 7DPA in two different data sets (Dean et al. 2011; Belmonte et al. 2013). **B,** Overlap of all four sets of genes. **C,** Genes that are more highly expressed tend to be near more accessible DHSs and vice versa. P-values are calculated using the hypergeometric test. One asterisk (*) indicates p-value < 0.01. Two asterisks (**) indicate p-value < 10^−20^.

For both data sets, genes with changing expression in seed coat between 4DPA and 7DPA stage were significantly more likely to reside near one or more dynamic DHSs (**Figure 2**). Furthermore, increased chromatin accessibility was significantly associated with increased expression levels at both the 4DPA and 7DPA stage (**Figure 2C**). Conversely, decreased chromatin accessibility was associated with decreased expression levels; however, this association was not always statistically significant (**Figure 2C**).

Although 4DPA seeds are mainly in the globular stage of development, some will have progressed to the heart stage (Le et al. 2010; Chen et al. 2015). The INTACT transgene promoter (GL2) is activated in the embryo of both heart (4-5 DPA) and linear cotyledon (7 DPA) stage seeds. Therefore, we also examined the relationship of dynamic DHSs with genes differentially expressed between the heart and linear cotyledon stage seeds in seed coat and embryo proper (**Supplemental Figure 2A, B**). As with the globular vs linear cotyledon stage comparison, differentially expressed genes in seed coats were significantly more likely to reside near one or more dynamic DHS (1.58-fold). Genes differentially expressed in embryo proper were somewhat less, albeit significantly, likely (1.17-fold) to reside near one or more dynamic DHS.

We next explored whether genes neighboring multiple dynamic DHSs were enriched in gene sets previously identified to be involved in seed coat development as well as in genes with differential expression in the aforementioned studies. Indeed, there was a monotonic increase in fold-enrichment for each of these three data sets when examining genes neighboring one or more, two or more, or three or more dynamic DHS (**Supplemental Figure 2C**). This tendency was particularly visible for the smaller set of 48 genes with known roles in seed development, pointing to the presence of multiple DHSs as support for possible functional relevance.

### Genes near dynamic DHSs are implicated seed coat biology

To test whether the genes that resided near dynamic DHSs were involved in known seed coat biology, we analyzed their GO terms using GOstats (**Figure 3; Supplemental Tables 7, 8**). Genes residing near deactivated DHSs were enriched for development, regulation, response, and pigment genes. Genes nearest to activated DHSs were enriched in genes related to transport, cell wall, biosynthetic process, and localization, consistent with the known developmental processes occurring at this stage and the annotations for the twenty known seed coat development genes that resided near activated DHSs (**Figure 3C)**.

**Figure 3.**
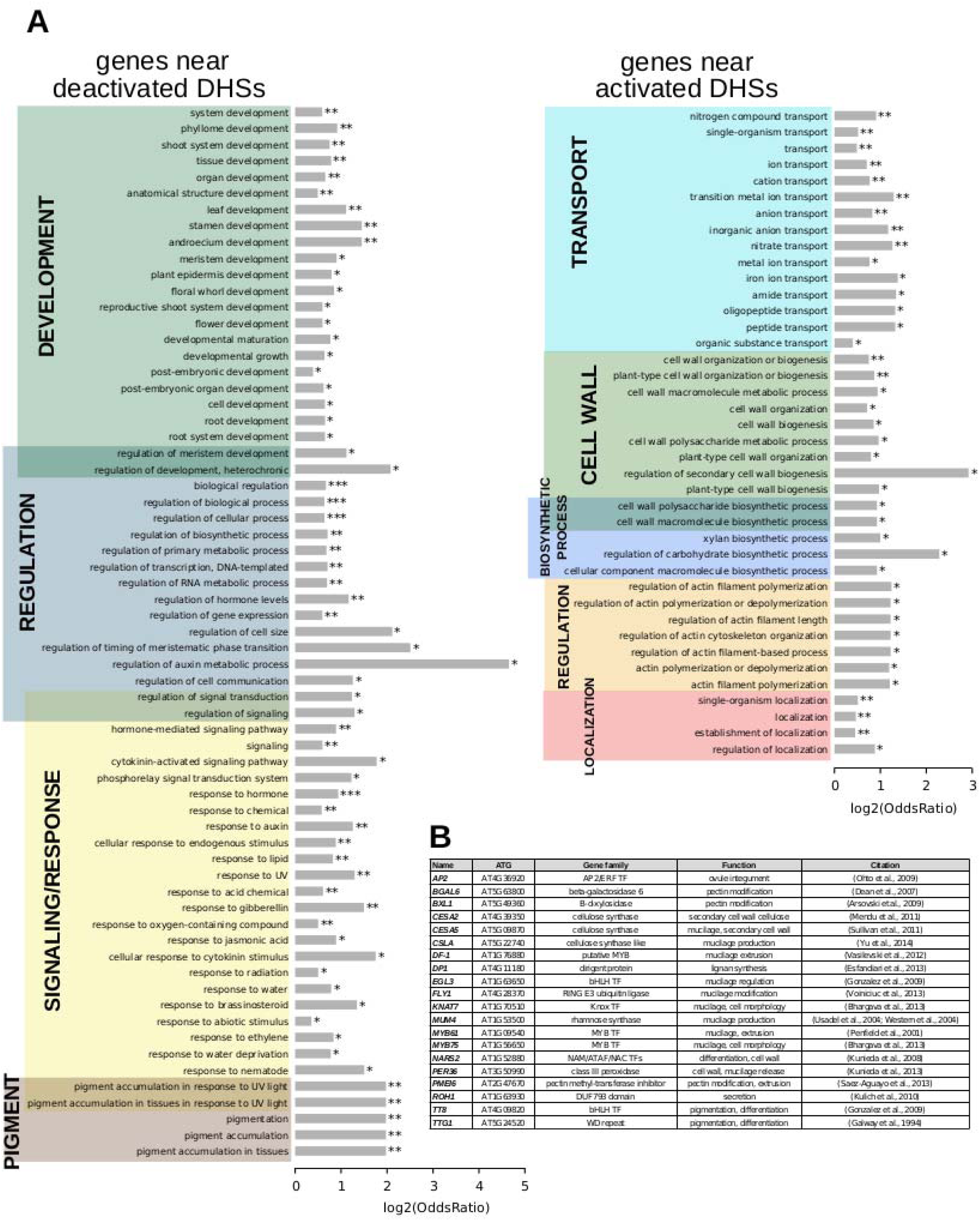
Term enrichment for genes nearest to dynamic DHSs. **A,** Term enrichment for genes near DHSs that are deactivated (less accessible, left) or activated (more accessible, right) at the 7DPA time point. **B,** Functional annotation for the twenty genes of the 48 known seed coat development genes that neighbor one or more dynamic DHS.

### Motif families in activated and deactivated DHSs are distinct

To determine candidate transcription factors driving dynamic DHSs in seed coat development, we examined transcription factor motif enrichments, comparing developmentally dynamic DHSs to union DHSs using AME (McLeay and Bailey 2010). Motifs for different TF families were enriched in activated versus deactivated DHSs compared to union DHSs. Specifically, bHLH and TCP motifs were significantly enriched in deactivated DHSs (**Figure 4A**). Motifs for many more transcription factor families were enriched in activated DHSs, including ARID, bZIP, MADS, MYB, MYB-related, and NAC motifs, with the majority of motifs belonging to either MYB and NAC transcription factors (**Figure 4B**). Previous functional studies validate our motif findings, lending support for novel associations of transcription factor motifs with seed coat development. For example, TCP3 overexpression leads to ovule integument growth defects and ovule abortion (Wei et al., 2015). In cotton, TCPs contribute to fiber elongation; cotton fibers like seed coat cells arise from the ovule outer integument. MYB61 is required for mucilage deposition and extrusion (Penfield et al., 2001), and NAM (NARS2) is a NAC TF important for differentiation and cell wall deposition (Kunieda et al., 2008).

**Figure 4.**
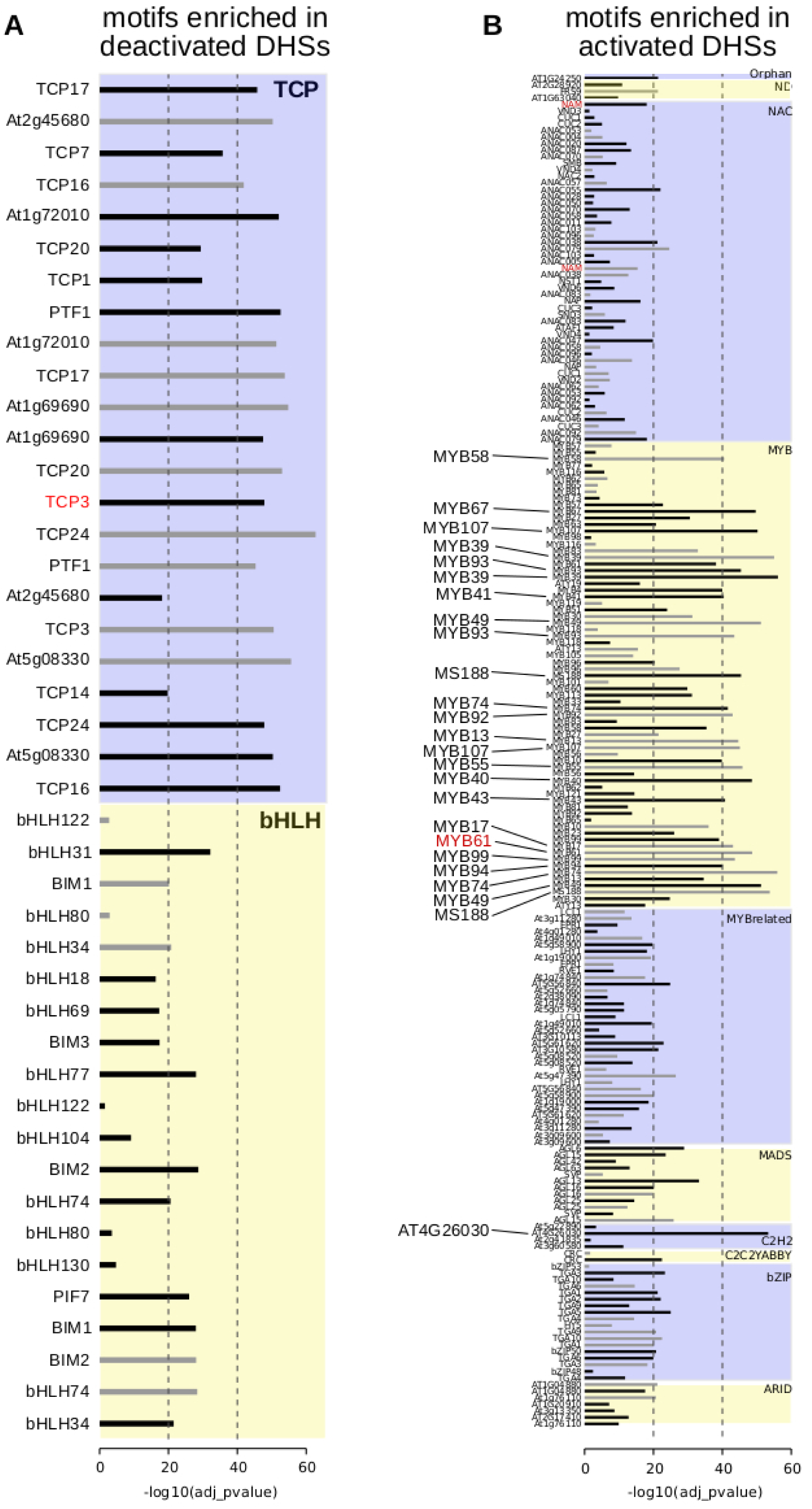
Motif enrichments within dynamic DHSs. **A,** Transcription factor motifs enriched in DHSs that are deactivated at the 7DPA time point. **B,** Transcription factor motifs enriched in DHSs that are activated at the 7DPA time point. Dotted vertical line indicates adjusted p-values of 10^−20^ of 10^−40^, respectively. All transcription factor family members are displayed if at least one member is enriched with adjusted p-value of 10^−20^ or less (greater than −log10(10^−20^) or 20). Transcription factor motifs derived using amplified (*i.e.*, non-methylated) DNA have gray bars indicating enrichment p-value (O’Malley et al. 2016). Motifs derived from genomic (*i.e.*, methylated) DNA have black bars indicating enrichment p-value.

### Comparative analysis of diverse plant regulatory landscapes

Previous studies in humans comparing regulatory landscapes of many cell types revealed cell lineage is encoded in the accessible regulatory landscape (Stergachis et al. 2013). Similarly, a dendrogram generated using accessibility profiles generated from thirteen diverse plant samples primarily reflected ontogeny; in contrast, treatment with major plant hormones and or severe stress mattered little for the regulatory landscape at large (**Figure 5A**). For example, the regulatory landscape of light-grown seven-day old seedlings inhabited a clade together with those of other light-grown seedlings that were either exposed to a severe heat shock or the plant hormone auxin. Both treatments are known to cause dramatic but drastically different changes in gene expression; yet, these did not suffice to obscure the commonalities in the regulatory landscapes of light-grown seedlings. Similarly, dark-grown seedlings, which differ profoundly in development from light-grown seedlings, clustered together. On a finer scale, the regulatory landscape of dark-grown seedlings exposed to the light-mimicking plant hormone brassinazole (BRZ) clustered closely with that of seedlings exposed to light for 24 hours before harvest, whereas the landscapes of seedlings exposed to shorter light treatments before harvest and seedling grown in the dark only were more distant. Overall, the regulatory landscapes of seedling tissue, both light and dark-grown were more similar to one another than those of the two epidermal cell types included in the analysis. The regulatory landscapes enriched for seed coat cells differed profoundly from those found in root hair and non-hair cells. This tendency is also evident in a Principal Component Analysis biplot, showing the sample vectors projected on the PC1-PC2 plane (**Figure 5B**). Our result are consistent with a meta study showing that expression profiles differ more among different tissues than among tissue-controlled treatments (Aceituno et al. 2008).

**Figure 5.**
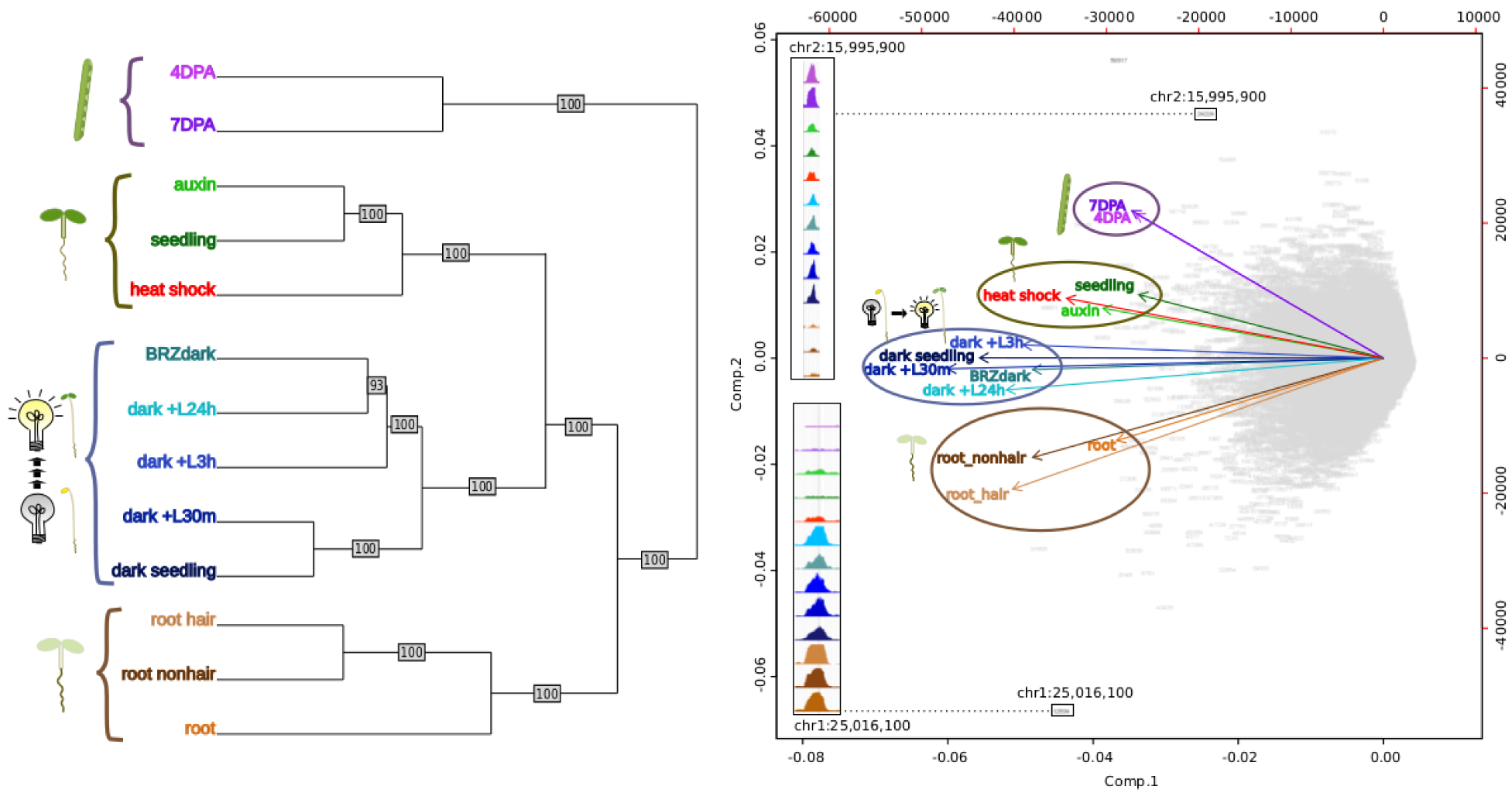
Comparative analysis of DHS landscapes in diverse samples. **A,** Dendrogram of thirteen samples using DNase I accessibility data. 4DPA, 7DPA denotes seed coat-enriched samples; auxin denotes 7-day-old seedlings treated with auxin (SRR8903039); seedling denotes 7-day-old control seedlings (A. M. Sullivan et al. 2014); heat shock denotes 7-day-old seedlings treated with heat shock (A. M. Sullivan et al. 2014); BRZ denotes 7-day-old seedlings treated with brassinazole (SRR8903038); dark+L24h, dark+L3h, dark+L30m denote 7-day-old seedlings which were grown in the dark and exposed to a long-day light cycle for the indicated amount of time, modeling development during photomorphogenesis (h, hours; m, minutes) (GSM1289351, GSM1289355, GSM1289353, respectively) (A. M. Sullivan et al. 2014); dark seedling denotes day-old dark grown seedlings (GSM1289357) (A. M. Sullivan et al. 2014); root hair denotes root hair cell samples of 7-day old seedlings (SRR8903037); root nonhair denotes nonhair root cells of 7-day-old seedlings (GSM1821072) (A. M. Sullivan et al. 2014); root denotes whole root tissue (GSM1289374) (A. M. Sullivan et al. 2014). **B,** Biplot of Principal Component Analysis of 62,729 DHSs by 13-sample matrix. Numbers in gray represent union DHSs. Insets show differential accessibility for two DHSs that were highly informative for distinguishing the 13 samples (*i.e.* these DHSs were among the most differentially accessible across all 13 samples). The upper inset shows a DHS that appears to be specific to aerial tissue; the lower inset shows a DHSs that appears to be specific to dark-grown tissue as roots are typically not exposed to light.

In animals and humans, each sampled cell type, tissue, or condition yields novel DHSs (Stergachis et al. 2013). Published studies in plants typically only sample a limited number of conditions or tissues, falling short of denoting comprehensive regulatory landscapes. We first determined which sample pairs yielded the most dynamic DHSs (**Figure 6**). Comparing the seed-coat enriched samples to one another yielded many more dynamic DHSs than any other comparison. The regulatory landscapes for the terminally differentiated root hair and root non-hair cells yielded the lowest number of dynamic DHSs.

**Figure 6.**
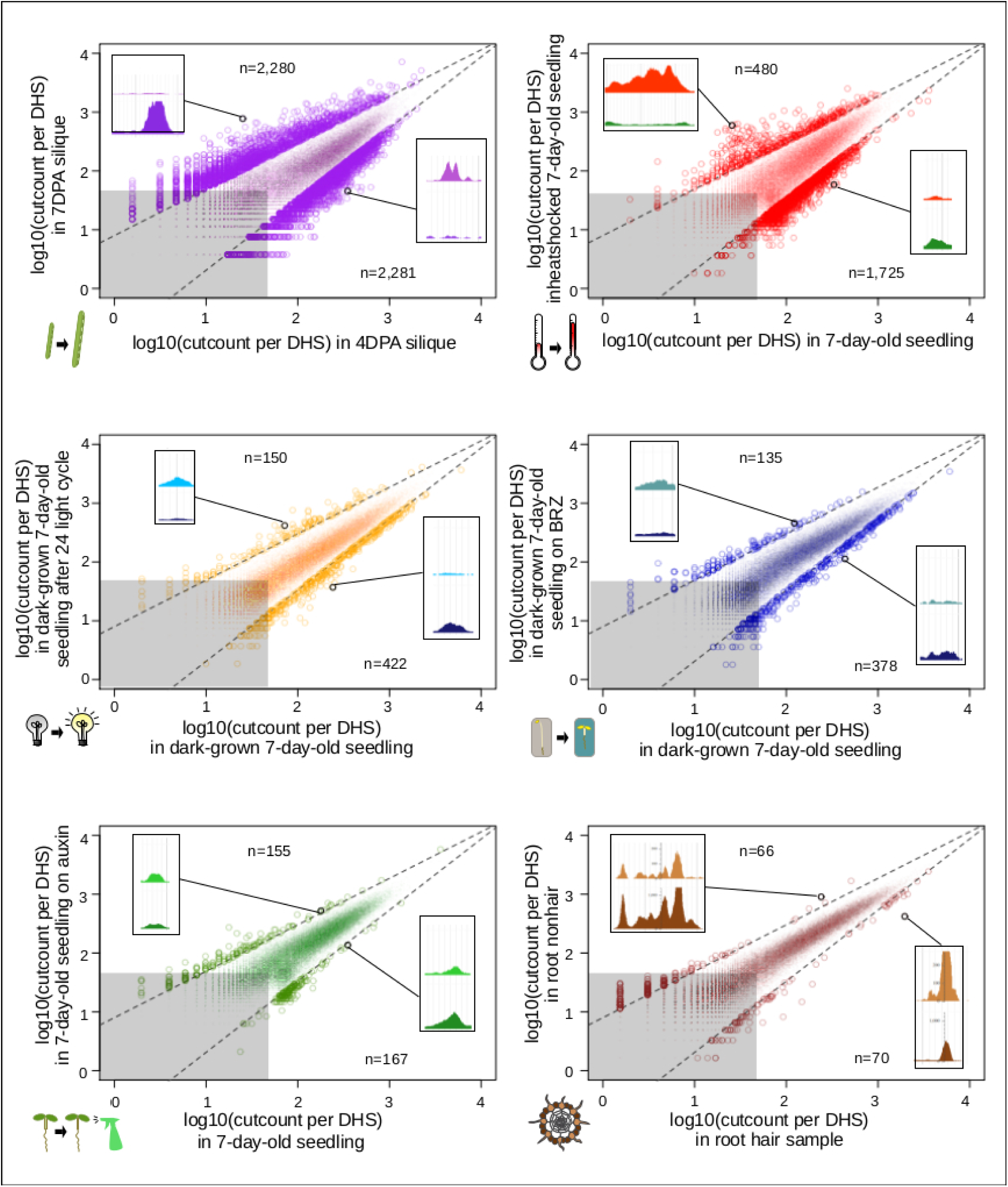
Comparison of seed coat-enriched samples (4DPA and 7DPA) results in the highest number of developmentally dynamic DHSs identified among all pairs examined. Scatterplots of log10(cut counts per union DHS) for six pairwise comparisons. Dotted lines creating a cone capturing the majority of the dots are drawn in the same location on each graph. Gray boxes represents regions in which both samples have less than 50 [*log10(50)=1.69897*] cleavage sites in that DHS. Numbers indicated above and below indicate the number of dots (DHSs) that lie above and below dotted lines. Screenshot insets in each graph showing an example differential DHSs above and below dotted lines are the following DHSs, respectively: {4DPA vs 7DPA: chr2:19,564,381-19,564,531, chr4:11,981,161-11,981,351; root hair vs root nonhair: chr1:30,035,761-30,036,071, chr4:280,861-281,131; control vs auxin-treated: chr1:10,320,801-10,321,131, chr1:5,204,361-5,204,551; dark-grown seedling vs dark-grown seedling on BRZ: chr5:22,570,821-22,571,231, chr5:21,869,241-21,869,591; control vs heat shocked seedling: chr4:7,338,681-7,342,041, chr2:18,374,201-18,374,371; dark-grown seedling vs dark-grown seedling exposed to 24hr light cycle: chr3:6,023,601-6,023,871, chr5:5,968,041-5,968,291}

For analyzing all 13 samples together, we merged their DHSs, excluding those below a certain cut count (marked in gray in **Figure 6**), thereby generating 46,891 union high-confidence DHSs, covering 10,374,430 bases or ∼7.4% of the genome (see **Methods** for details). We then excluded each of the thirteen samples individually, assessing how many hypersensitive bases unique to the sample were lost. The seed coat-enriched samples (both 4DPA and 7DPA) contributed the most sample-specific hypersensitive bases, followed by those found in whole roots (**Figure 7A**). Of the hypersensitive bases identified in the seed-coat-enriched samples, over half (2,858,990 bps / 5,573,620 bps) were not present in 7-day-old light-grown seedlings, and over 25% (1,418,070 bps / 5,573,620 bps) were not present in any of the other eleven samples examined. As more and more samples are tested, the number of identified hypersensitive base pairs is expected to plateau. We observe this phenomenon already with the 13 samples included (**Figure 7B**). Note, however, that our analysis underestimates overall DHS frequency due to subsampling all samples to the lowest read-coverage sample (14 million reads, see **Methods**). Increasing read coverage increases the number of identified hypersensitive base pairs up to a saturation point, which depends on genome size. For the small genomes of *A. thaliana* and *D. melanogaster*, this saturation point is reached with ∼20 million reads for a given sample; using 14 million reads will identify ∼70% of the DHSs identified with 20 million reads.

**Figure 7.**
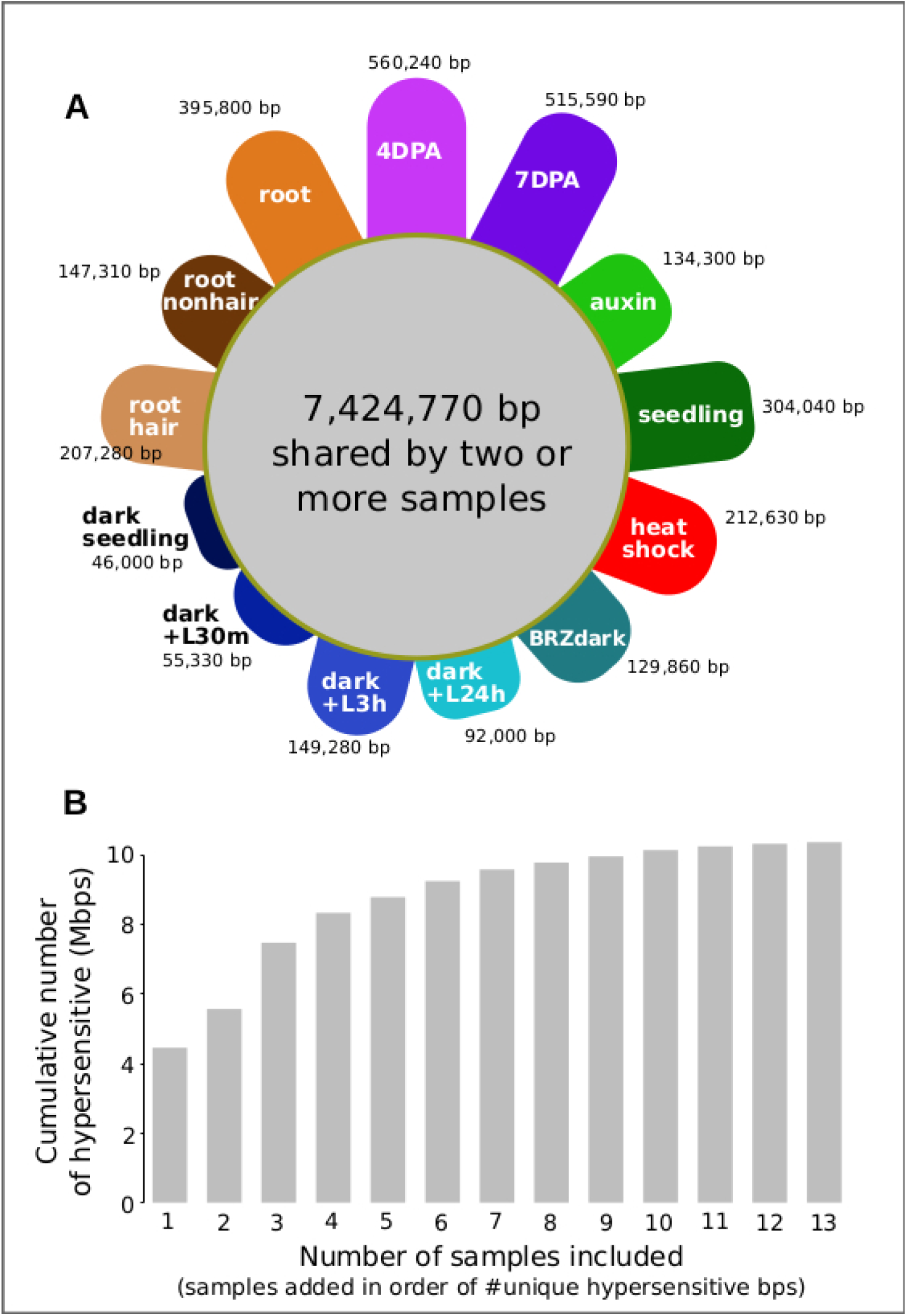
Seed coat-enriched samples contribute the largest number of novel hypersensitive bases in a diverse set of samples. **A,** Colored petals denote number of unique hypersensitive base pairs in each sample, gray circle denoted hypersensitive base pairs shared by two or more samples. Sample labels as in Figure 5; samples are grouped by seed coat-enriched samples, light-grown seedlings, dark-grown seedlings, and root samples. **B,** Cumulative number of hypersensitive sites plateaus. Graph was generated by adding samples based on their number of unique hypersensitive base pairs, starting with the largest (4DPA) and ending with the smallest (dark seedling).

## Discussion

Here, we mapped regulatory elements and their developmental dynamics in *GL2*-expressing cells from whole siliques using DNase I-seq. We targeted the developmental stages in which the seed coat transitions from a state of growth to a state of mucous production and secretion. During this developmental window, more than 3,000 DHSs changed reproducibly in accessibility.

DHSs are a hallmark of regulatory DNA and thus dynamic DHSs often reside in close proximity to genes with changing expression. However, it is well-established that the association between chromatin accessibility, even if dynamic, and nearby gene expression is imperfect for several reasons (A. M. Sullivan et al. 2015). First, regulatory DNA is often be poised, *i.e.* bound by transcription factors and hence accessible, without transcription occurring (Elgin, 1988); in addition, DHSs often remain accessible after transcription has occurred (Groudine and Weintraub, 1982). Second, the binding of both activators (Morgan et al., 1987) *and* repressors (Baniahmad et al., 1990) can remodel chromatin locally causing increased accessibility. Therefore, increases in chromatin accessibility do not necessarily translate into increases in gene expression. Finally, distal regulatory elements, *i.e.* enhancers residing in intergenic regions, can function at long distances and are agnostic to orientation (Banerji et al., 1981). Compared to union DHSs, we found that more dynamic, differentially accessible DHSs in seed coat-enriched cells resided in intergenic regions. As we assigned DHSs to target genes based on proximity, we will have missed long-range interactions, possibly assigning incorrect target genes. Nevertheless, we observed considerable agreement between the direction of changes in chromatin accessibility and changes in expression for neighboring genes.

Despite these limitations, dynamic DHSs are potentially useful for identifying new candidate genes that control seed coat development; moreover, their motif enrichments can point to the TFs that drive the observed DHS and gene expression dynamics. Genes near deactivated DHSs (up in 4DPA) were associated with development, signaling, pigment, and regulation, consistent with the processes occurring during seed maturation. Genes near activated DHSs (up in 7DPA) were associated with secretion, localization, biosynthetic processes, and cell wall modification, consistent with these cells switching to mucous production and secretion into the apoplast, and ramping up to build the columella, a secondary cell wall structure. Although most differentially expressed genes resided in close proximity to only one dynamic DHS, several hundred genes neighbored as many as ten dynamic DHSs, consistent with multiple regulatory inputs during development. Genes neighboring multiple dynamic DHSs were enriched for genes with altered expression in seed coat development. This trend was most strongly observed in known seed coat development genes. We have noted previously that genes conditionally expressed in response to abiotic treatments tend to neighbor multiple DHSs (Alexandre et al. 2017). It appears that multiple DHSs are also a feature of developmentally dynamic genes.

Motif enrichments within activated and deactivated DHSs revealed distinct transcription factor families and individual transcription factors that may be regulating seed coat maturation. Among the TF motifs most enriched in deactivated DHSs were those of the TCP family. TCPs are involved in many aspects of development, particularly in land plants in which the class has greatly diversified (Martín-Trillo and Cubas 2010). Consistent with its significant motif enrichment in deactivated DHSs, overexpression of *TCP3* leads to ovule integument growth defects and ovule abortion in *A. thaliana* (Wei et al., 2015). Altered expression of the most famous member of the TCP TF family, the maize TF *tb1*, contributes to the morphological changes in shoot architecture that differentiate wild teosinte and domesticated maize (Clark et al., 2006).

Among TF motifs most enriched in 7DPA-activated DHSs were those of the MYB family. This class of TFs, present throughout Eukarya, plays important roles in plant development and stress responses (Ambawat et al. 2013). All of the MYB TFs with enriched motifs in activated DHSs belonged to the same subfamily, the R2R3 MYBs, which are involved in secondary metabolism and cell fate establishment (Stracke, Werber, and Weisshaar 2001). MYB61, whose motif is enriched in our analysis, is required for mucilage production and secretion in cell coat cells (Penfield et al., 2001). Zinc finger, MADS-box, and AT-hook TFs were also enriched in 7DPA-activated DHSs; these TF families have not been implicated previously in seed coat cell maturation. However, MADS-box TFs are required for proper ovule development (Honma and Goto 2001; Pinyopich et al. 2003).

This foray into cell-type-specific regulatory landscapes in plants, an approach that has been previously pioneered in humans and animal models and indeed has been the primary mode of analysis in these systems demonstrates the dramatic coverage and knowledge gains by analyzing specific cell types and their developmental dynamics rather than using whole seedlings or easily dissected tissues. Specifically, a single whole seedling sample previously yielded 34,288 DHSs covering ∼4% of the *A. thaliana* genome (A. M. Sullivan et al. 2014). Our combined analysis of seed coat cells and 11 other samples generated a set of 46,891 union DHSs which accounted for ∼7.4% of the *A. thaliana* genome. Of these, 1,978 were entirely non-overlapping with DHSs in the other 11 samples. Expressed in base pairs this result appears even more impressive: of 10,374,430 hypersensitive, accessible bps in all 13 samples, 560,240 hypersensitive bps (>5%) were unique to the seed coat-enriched samples. This result demonstrates that cell-type-specific DHS profiling holds enormous promise for expanding our knowledge of the *A. thaliana* regulatory landscape. Although heat stress, auxin, and brassinazole treatments cause dramatic changes in genes expression, our comparative analysis shows that cell lineage and developmental stage rather these treatments are reflected in regulatory landscapes, which is consistent with prior knowledge of poised transcription factors (Elgin, 1988), in particular those occupying heat shock promoters (Vihervaara, Duarte, and Lis 2018). Our findings argue for exploring regulatory landscapes across all plant cell types, across development, and in response to relevant conditions to fully understand understand how chromatin accessibility and gene expression are integrated into precise expression patterns. The regulatory elements identified in this study can now be integrated with the existing co-expression- and genetics-based gene regulatory network data to gain a more complete understanding of the regulation of seed coat maturation (Francoz et al., 2015).

## Acknowledgements

This work was supported by grants from the National Science Foundation (MCB1243627 to JS, CQ, JN, MCB1516701 to CQ, and NSF RESEARCH-PGRP 1748843 to CQ), and Graduate Research Fellowship (DGE-0718124) (AMS). We thank Roger Deal and Steven Henikoff for sharing INTACT lines and experimental expertise, members of the Stamatoyannopoulos and Queitsch labs for useful discussions. We thank Chris Gee for technical assistance and G. Alex Mason for drawing and colorizing the seeds at different stages of development. The authors have no conflicts of interest.

## Author Contributions

A.A.A. designed the experiments, and together with A.T. executed DNase I-seq experiments. A.M.S. and K.L.B performed data analysis and made the figures. K.L.B., R.S, R.E.T, S.N., A.K.J. and S.T.S assisted with the bioinformatic analysis and data processing. P.J.S, F.V.N, M.W and M.D. sequenced DNase I-treated samples. A.M.S., K.L.B. and C.Q. wrote the manuscript. J.A.S. and J.L.N facilitated experiments and assisted in writing the manuscript. All authors read, commented on, and approved the manuscript.

## Methods

### Sample preparation

Siliques of appropriate ages from the INTACT line *GL2_pro_:NTF/ACT2_pro_:BirA* (Deal and Henikoff 2010) were collected by first marking young flowers using a fine paint brush and water based paint as previously described (Western, Skinner, and Haughn 2000). In brief, recently opened flowers are chosen at the stage the anthers are almost at the same level as the pistil and fertilization is able to occur, usually 2 per plant per day at this stage. The flower is marked with paint and silique collected 4 or 7 days later. Samples were prepared using INTACT nuclei isolation (Deal and Henikoff 2010) followed by DNase I-seq (A. M. Sullivan et al. 2014). A detailed protocol for tissue preparation and nuclei isolation using INTACT lines is provided at plantregulome.org. A detailed protocol for post-digestion sample processing has been published previously (John et al. 2013). Data sets may be found in GEO accessions GSE53322 and GSE53324 and at plantregulome.org.

### Microscopy

#### Testing activity of the INTACT construct in seed coat cells

Whole seeds were observed on a Leica TCS SP5 II laser scanning confocal microscope. Whole seed images (**Supplemental Figure 1A**) are z-stack composites of 35 individual images using an HC Plan Apo CS 20X objective. Image of seed coat cell layer (**Supplemental Figure 1B)** is a single image using the 63X water immersion objective.

### Data processing for seed coat analysis

Five DNase I-seq libraries, including biological replicates for each time point, were sequenced and aligned to the TAIR10 reference genome using bwa/0.6.2. Because number of peaks called is a function of read depth, 24 million reads mapping to chromosomes 1-5, excluding centromeres (chr1:13,698,788-15,897,560; chr2: 2,450,003-5,500,000; chr3:11,298,763-14,289,014; chr4:1,800,002-5,150,000; chr5:10,999,996-13,332,770), were sampled from the biological replicate with the highest read coverage for each developmental time point (4DPA-DS20201 and 7DPA-DS21306). These 24M-read bam files were used to call DHSs (peaks) using the HOTSPOT program (John et al. 2011a). DHSs from these two samples were merged to create a union set of 43,120 DHSs. DESeq2 (Love, Huber, and Anders 2014) was used on this set of union DHSs to identify a subset of 3,440 developmentally dynamic DHSs (adjusted p-value < 0.01), using all reads mapping to chromosomes 1-5, excluding centromeres, from all five samples (4DPA-DS20201, 4DPA-DS20131, 4DPA-DS20132, 7DPA-DS21306, 7DPA-DS20134). We then removed DHSs with mean cut count of 50 or less -- roughly the bottom ten percentile -- leaving 3,109 dynamic DHSs. Data sets may be found in GEO accessions GSE53322 and GSE53324 and at plantregulome.org.

### Genomic distribution of DHSs

DHS midpoints were used to determine overlaps with genomic elements. Genomic elements (5’UTR, coding regions, 3’UTR, intergenic, TE) were extracted from the TAIR10 gff file on arabidopsis.org. Centromeric regions were excluded from the analysis. To simplify the analysis, only the primary transcript of each gene (AT*.1) was considered. When a single DHS midpoint coincided with two different elements, both element overlaps were tallied, thus overlapping DHS counts sum to greater than the initial number of DHSs. We tallied the total number of base pairs within each element type in the genome, double-counting base pairs that are assigned to overlapping elements. Tallies may be found in **Supplemental Table 6**.

### Integration with expression data sets

Genes from Dean et al. 2011 and Belmonte et al. 2013 were considered to be differentially expressed if there was a 2-fold change in expression between time points. Dean et al. 2011 identify the genes that change 2-fold between 3DPA and 7DPA; these genes were used for integration with dynamic DHS data. The genes that change expression by 2 or more fold in Belmonte et al. 2013 were extracted from the published normalized expression data (**Dataset S2**). We used the hypergeometric test to measure how different the observed number of DHS-gene pairs in certain configurations were compared to the expected number. For example, there were 2,131 genes that had 2-fold more expression at 7DPA than 3-4DPA in the Belmonte et al. 2013 data set, and 3,269 genes that were near dDHSs that were more accessible at 7DPA than 4DPA. Given that there are 28,775 genes total, we expect 2,131 x 3,269 / 28,775 ≈ 242 DHS-gene pairs with this configuration if accessibility and expression are randomly associated. We observe 586 such DHS-gene pairs, which is a statistically significant excess (p-value < 10^−20^).

### Term enrichment

Term enrichments were performed using the org.At.tair.db (Carlson 2016) and GOstats (Falcon and Gentleman 2007). Only the enrichments with a p-value less than 0.001 are shown in **Figure 3**.

### Motif enrichment

Enrichment of motifs (O’Malley et al. 2016) in sequence underlying dDHSs as compared to union DHSs was evaluated using AME (McLeay and Bailey 2010). All members of motif families in which at least one member is enriched with significance of p<10^−20^ are displayed in **Figure 4**. All motifs with corrected p-value of less than 0.01 are listed in **Supplemental Tables 9&10**. Motifs derived using amplified DNA (colamp_a) are gray and motifs derived using native genomic DNA (col_a) are black.

### Comparative analysis of DHS landscapes

Each of 13 samples was subsampled to roughly 14 million reads mapping to chromosomes 1-5, excluding centromeres (chr1:13,698,788-15,897,560; chr2: 2,450,003-5,500,000; chr3:11,298,763-14,289,014; chr4:1,800,002-5,150,000, chr5:10,999,996-13,332,770) (**Supplemental Table 11)**. DHSs were called on these 13 bam files using the HOTSPOT program (John et al. 2011b), and a union set of DHSs was generated by merging DHSs from each of these 13 samples with BEDOPS (Neph et al. 2012), (bedops –m, adding each sample in succession) (**Supplemental Table 12**). There were 62,738 DHSs in this union set. Per-base DNase I cleavages (cut counts) within each union DHS were tallied for each sample. Cleavage tallies were normalized for sample quality by dividing by the proportion of DNase I cleavages within 1% FDR threshold hotspots.

#### Accessibility profiles used to cluster samples

Dendrogram and bootstrap values were generated 100 trees from random subsamples of 10,000 DHSs using the ape package (Paradis, Claude, and Strimmer 2004). Principal Component Analysis was performed on the 62,729 by 13 matrix. For the PCA, we excluded nine DHSs within the first 50 kb of chromosome 2, part of a NOR (nucleolar organizer region) (Copenhaver and Pikaard 1996; Lin et al. 1999), a region with unusually high cut count, similar to the centromeres.

#### Sample-specific hypersensitive bases

To identify sample-specific hypersensitive bases, we merged large DHSs (>50 cleavages per DHS) from the 13 samples to generate a set of 46,891 union DHS covering 10,374,430 bps. We then generated 13 new merged sets of DHSs using only 12 samples, excluding one of the samples in each set, and then determined the number of hypersensitive bases not captured. We define the number of hypersensitive bps unique to the sample as number of bps in the 13-sample union DHS set minus the number of bp in the 12-sample union DHS set divided by the number of bps in the 13-sample union DHS set (**Figure 7**).

#### Pairs of samples resulting in differential DHSs

To compare the number of developmentally dynamic DHSs identified with different pairs of samples, we used the complete set of merged DHSs (62,738 unionpeaks). For each of six pairwise comparisons, we made a scatterplot of the cut counts of these 62,738 unionpeaks. We then defined developmentally dynamic DHSs as those that both lie outside a cone defined by the lines *y=(1-0.21)x + 0.9* and *y=(1+0.21)x − 0.9* and have greater than 50 cleavages per unionpeak in at least one sample. Expression differences between these pairs have been previously published (A. M. Sullivan et al. 2014).

### Data Access

All DNase I-seq data are available at GEO (https://www.ncbi.nlm.nih.gov/geo/) and/or SRA (https://www.ncbi.nlm.nih.gov/sra/). 4DPA-DS20201: SRR5873456; 4DPA-DS20131: SRR5873454; 4DPA-DS20132: SRR5873455; 7DPA-DS21306: SRR5873453 and 7DPA-DS20134: SRR5873452). Auxin samples: SRR8903039. Seedling control sample: DS19992 GSM1289363. Heat shock sample: GSM1289361. BRZ sample: SRR8903038. Photomorphogenesis series samples: dark-DS22138 (GSM1289357), dark+L30m (GSM1289353), dark+L3h (GSM1289355), dark+ L24h (GSM1289351). Hair samples (root hair): SRR8903037. Nonhair sample (root nonhair): GSM1821072. Root sample: GSM1289374.

---

**Supplemental Figure 1.**
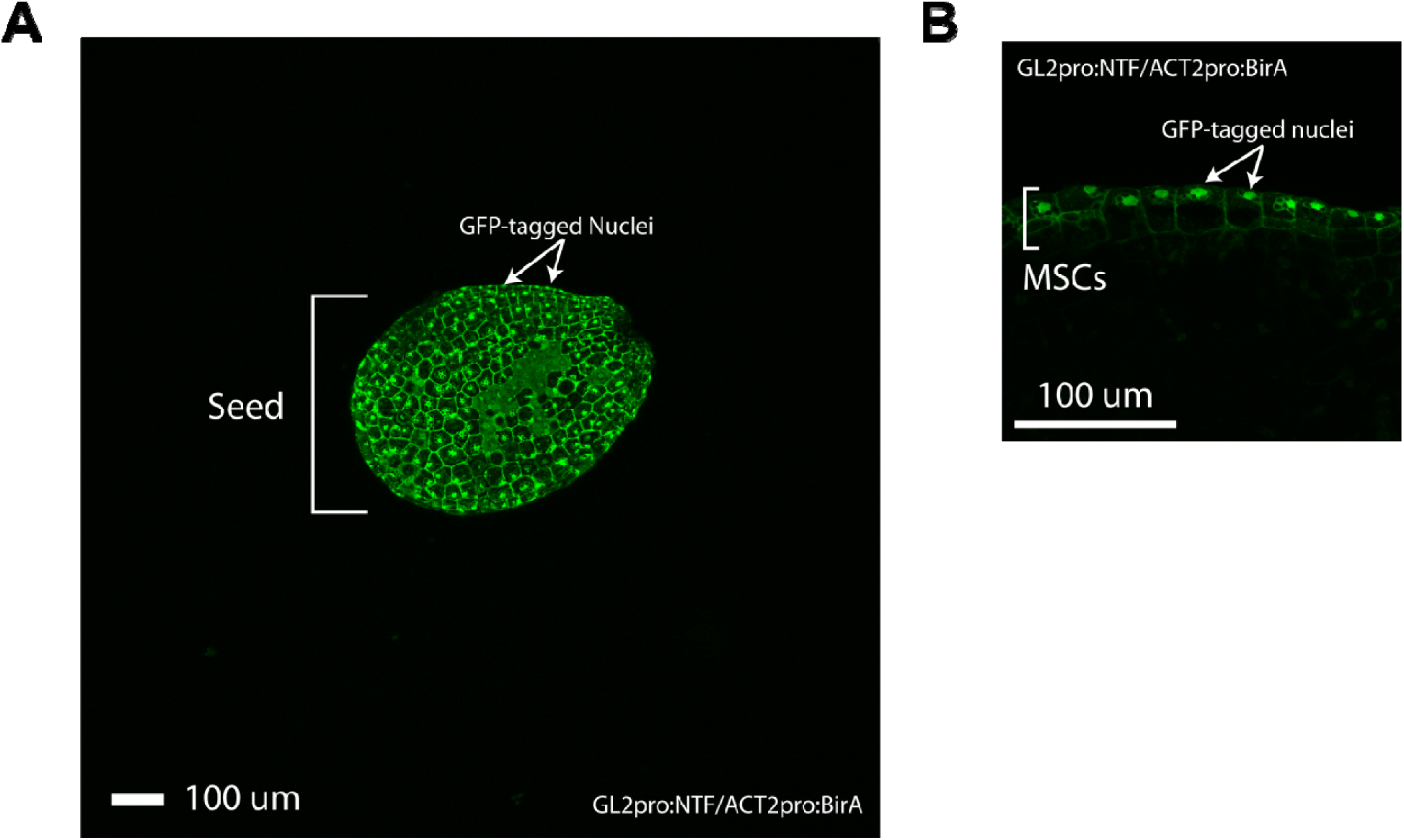
Confocal microscopy of INTACT-tagged nuclei in seed coat epidermis. **A.** Confocal of whole seed at 4DPA from the INTACT line *GL2_pro_:NTF/ACT2_pro_:BirA* (Deal and Henikoff, 2010). GFP-fluorescing nuclei are evident across the seed coat epidermis. Scale is 100um. **B.** Confocal of 4DPA mucous secreting cells (MSCs) from the INTACT line *GL2_pro_:NTF/ACT2_pro_:BirA* (Deal and Henikoff, 2010). GFP-fluorescing nuclei are readily observable in the outer most layer of the seed coat. Scale is 100um.

**Supplemental Figure 2.**
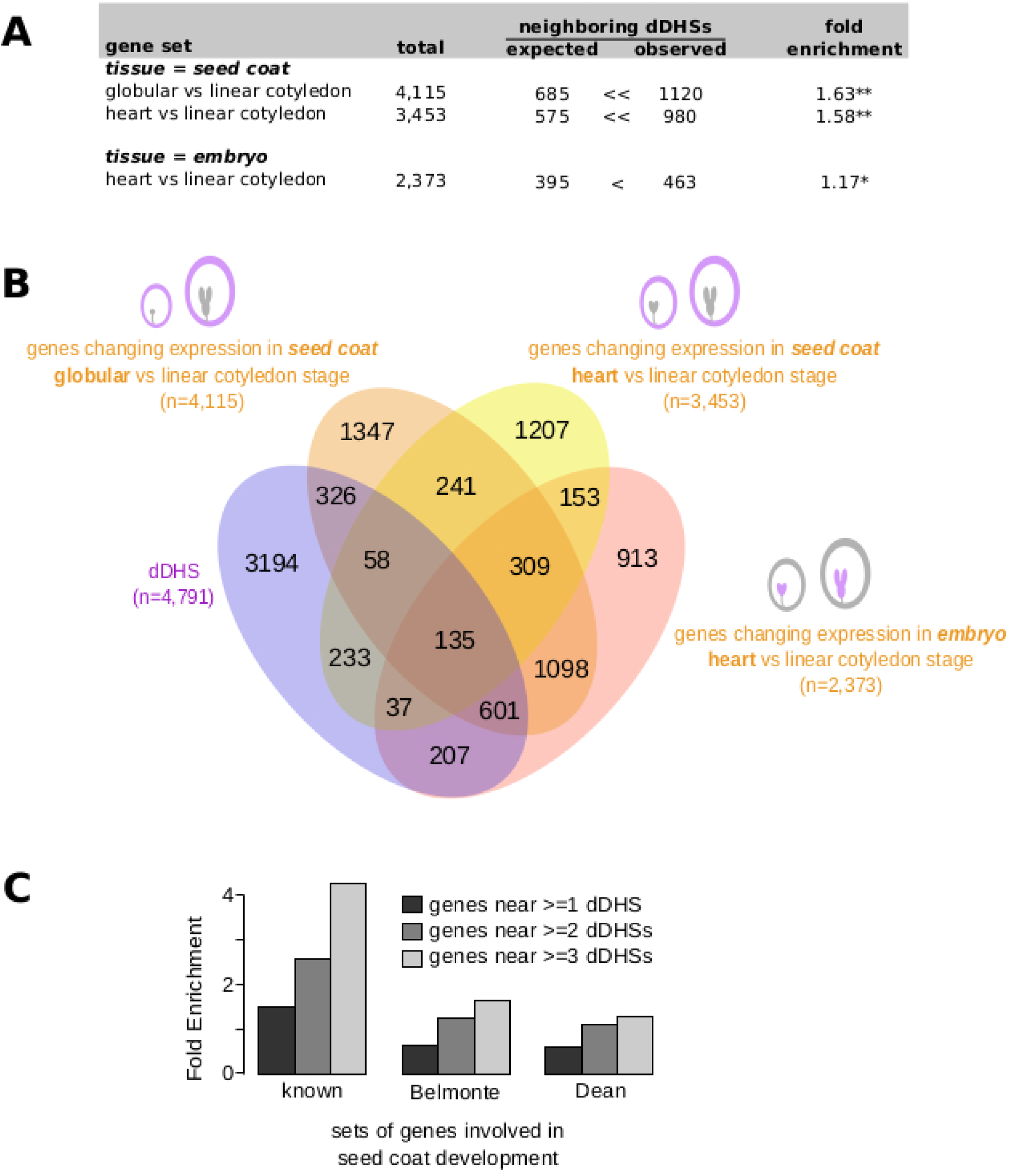
Genes neighboring developmentally dynamic DHSs are often differentially expressed in seed coat and embryo. **A,** Overlap between the set of genes neighboring dDHSs and genes differentially expressed in seed coat at globular vs linear cotyledon stage and heart vs linear cotyledon stage, and genes differentially expressed in embryo at heart vs linear cotyledon stage (Belmonte et al. 2013). One asterisk (*) indicates p-value < 0.01. Two asterisks (**) indicate p-value < 10^−20^. **B,** Overlap of all four sets of genes. **C,** Genes neighboring multiple dynamic DHSs tend to be more enriched for seed coat development genes. This is seen in the set of 48 known seed coat development genes (**Supplemental Table 1)** as well as in genes with differential expression (Dean et al. 2011; Belmonte et al. 2013).

